# FANCD2 Alleviates Physiologic Replication Stress in Fetal Liver HSC

**DOI:** 10.1101/2020.09.30.320796

**Authors:** Makiko Mochizuki-Kashio, Young me Yoon, Theresa Menna, Markus Grompe, Peter Kurre

**Affiliations:** Department of Pediatrics, Papé Family Pediatric Research Institute, Pediatric Blood & Cancer Biology Program, Stem Cell Center, Oregon Health & Science University, Portland, OR; Committee on Immunology, Graduate Program in Biosciences, University of Chicago, Chicago, IL; Department of Microscopic and Developmental Anatomy, Tokyo Women’s Medical University, Tokyo; Comprehensive Bone Marrow Failure Center, Children’s Hospital of Philadelphia; Perelman School of Medicine, University of Pennsylvania, Philadelphia, PA

## Abstract

Bone marrow failure (BMF) in Fanconi Anemia (FA) results from exhaustion of hematopoietic stem cells (HSC), but the physiological role of FA proteins in HSC pool integrity remains unknown. Herein we demonstrate that FANCD2, a core component of the FA pathway, counters replication stress during developmental HSC expansion in the fetal liver (FL). Rapid rates of proliferation and FANCD2 deficient result in excess RPA-coated ssDNA, and provoke pChk1 activation and *Cdkn1a(p21)* nuclear localization in fetal *Fancd2*^−/−^ HSC. Checkpoint mediated S-phase delays induced by *Cdkn1a(p21)* are rescued by Tgf-*β* inhibition, but pChk1 activation is further aggravated. Our observations reveal the mechanism and physiological context by which FANCD2 safeguards HSC pool formation during development.

## INTRODUCTION

Hematopoietic stem cells (HSC) are defined by their ability to self-renew and differentiate, whereby successive rounds of cell division give rise to increasingly specialized progenitor cells while maintaining a pool of multipotent HSC. To generate adequate regenerative capacity, fetal stem cells successively colonize different microenvironments, each providing unique cues for the formation of a finite pool of HSC clones that provide the basis for a lifelong supply of blood and immune cells (Mikkola and Orkin, 2006). The fetal liver (FL) takes on a crucial role for rapid clonal expansion during development. Not surprisingly, HSC rely on intact cell cycle checkpoints and DNA repair pathways to minimize the potential mutational burden in a highly proliferative HSC pool (Beerman et al., 2014; Schuler et al., 2019).

Fanconi Anemia (FA) is a cancer predisposition syndrome, and bone marrow (BM) failure early in life is a principal source of morbidity and mortality (Ceccaldi et al., 2012; Rosenberg et al., 2004). Compound heterozygous mutations in one of 22 FA genes that cooperate in a DNA repair pathway also lead to HSC deficits and symptomatic cytopenias by early school age. Indeed, even the youngest patients reveal depleted hematopoietic stem and progenitor cell (HSPC) populations (Ceccaldi et al., 2012; Kelly et al., 2007). Whereas murine models of FA mimic the *postnatal* p53-dependent, apoptotic HSC loss seen in patients only under experimental stress, we and others observed spontaneous deficits in the HSC pool of *Fancc*^−/−^, *Fancd2*^−/−^, *Fancg*^−/−^ fetuses (Botthof et al., 2017; Ceccaldi et al., 2012; Domenech et al., 2018; Kamimae-Lanning et al., 2013; Suzuki et al., 2016; Yoon et al., 2016). Neither the specific mechanism, nor physiologic stage of onset for hematopoietic failure in FA are currently known. Hematopoietic reserve in the adult is typically guarded by maintaining a majority of HSC in quiescence and successively activating individual clones to match demand. Accordingly, the formation of a sufficient pool of HSC to last a lifetime is tightly regulated, and any deficits in generating sufficient clonal diversity *in utero* will exert a disproportionate influence on the pace of postnatal hematopoiesis exhaustion.

Previously, several groups showed that FANCD2 is upregulated in response to experimental replication stress conditions (Balcerek et al., 2018; Chaudhury et al., 2013; Lossaint et al., 2013; Schlacher et al., 2011; Schlacher et al., 2012; Thompson et al., 2017; Tian et al., 2017). Here we show that this matches the physiological role in the FL without experimental provocation, making FANCD2 a critical component for HSC pool formation during development.

## RESULTS

### The fetal liver specifies a developmental window of vulnerability that governs FANCD2^−/−^ HSC losses

Steady state function and regenerative capacity of the hematopoietic system rely on a finite number of HSC, formed during development and sufficient to assure lifelong function (Mikkola and Orkin, 2006). To understand the origin of developmental deficits in FA, we first performed detailed profiling of HSPC subsets in WT and *Fancd2*^−/−^ mice at seven time points across ontogeny, in FL (E12.5, E13.5, E14.5 and E18.5), fetal BM at E18.5, postnatal BM at P21 and adult BM at 10- and 30-weeks. Results reveal near equivalent HSC (Lin^−^/Sca-1^+^/c-kit^+^/CD150^+^/CD48^−^) frequency at E12.5, with significant differences between the two genotypes first emerging during the ensuing expansion in the FL (**Fig. 1A**). The data are consistent with studies in *Fanca*^−/−^ mice (Kaschutnig et al., 2015), where differences in BM HSC frequency normalize rapidly postnatally and become non-significant after the P21 time point, when HSC assume a more quiescent adult phenotype (Copley et al., 2013). Analysis of myeloid-committed multipotent progenitors (MPP) 2 (Lin^−^/Sca-1^+^/c-kit^+^/CD150^+^/CD48^+^) (**Fig. S1A**) and the lymphoid progenitors enriched MPP3/4 population (Lin^−^ /Sca-1^+^/c-kit^+^/CD150^−^/CD48^+^) (**Fig. S1B**) indicate that *Fancd2*^−/−^ deficits give way to a myeloid predominant hematopoietic system, a known response to proliferative stress in the aging hematopoietic system (Pietras et al., 2015).

**Figure 1.**
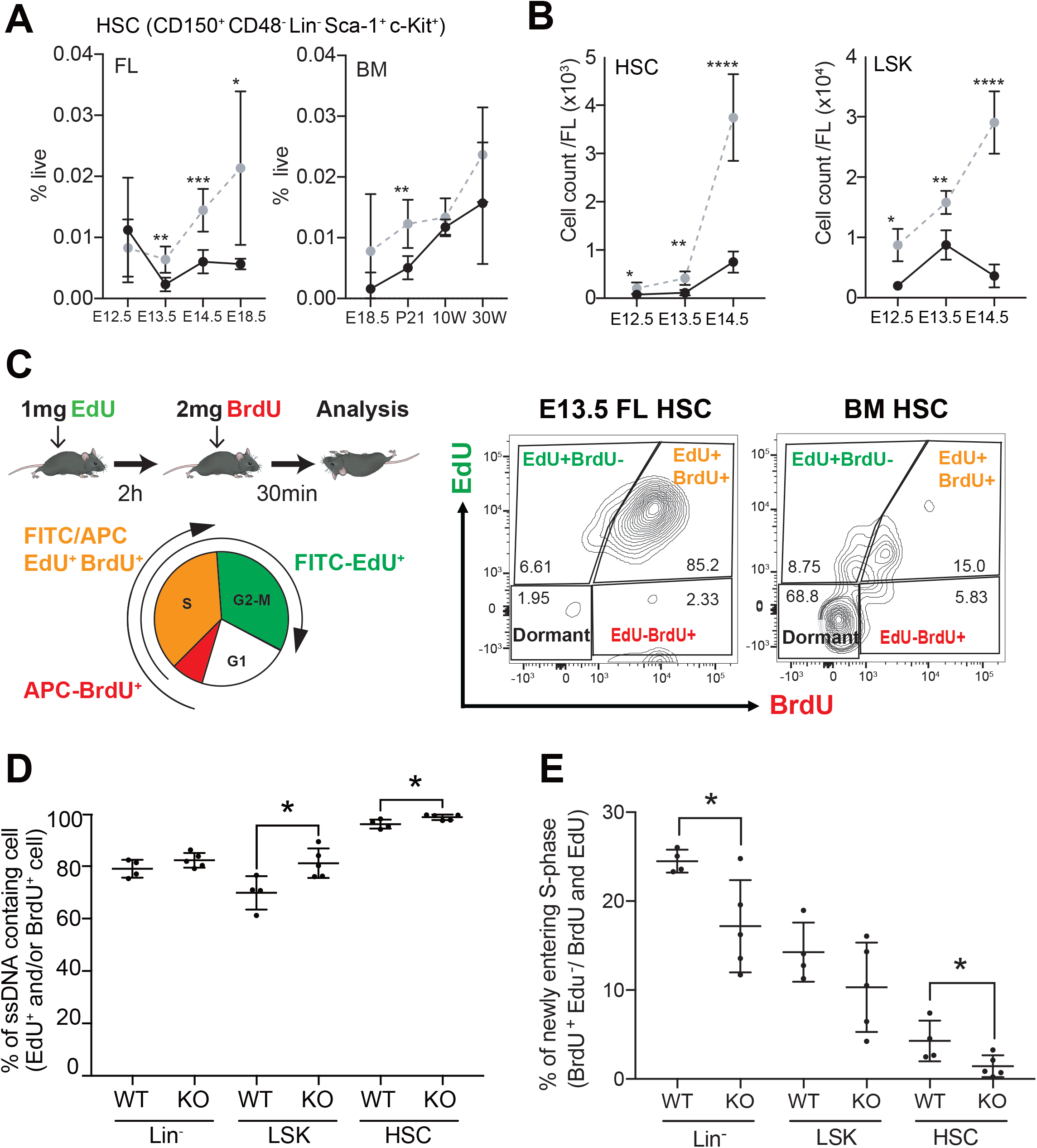
S-phase delay in Fancd2^−/−^ FL HSC. **(A,B)** Immunophenotyping was performed to determine the frequency of **(A)** CD150^+^ CD48^−^ Lin^−^ Sca-1^+^ c-Kit^+^ (LSK), HSC **i**n WT and Fancd2^−/−^ cell across select time points in ontogeny. **(B)** Absolute number of HSC (left panel) and LSK (right panel) in E12.5-13.5-14.5 of WT compared with Fancd2^−/−^ FL. **A,B:** E12.5 n=3^+/+^, 7^−/−^, E13.5 n=4^+/+^, 5^−/−^, E14.5 n=9^+/+^, 6^−/−^, E18.5 n=7^+/+^, 3^−/−^, P21 n=5^+/+^, 6^−/−^, 10 Weeks(10W) n=4^+/+^, 4^−/−^, 30W n=3^+/+^,3^−/−^. \**P*<0.05, \*\**P*<0.01, \*\*\**P*<0.001, \*\*\*\**P*<0.0001. **(C)** To understand midgestational deficits in HSC expansion, we performed kinetic cell cycles studies adapting a sequential EdU/BrdU injection protocol. Schema for sequential EdU/BrdU injection in the dam at E13.5 with cell cycle analysis. Representative flow panels illustrate FL (left) and BM (right) HSC distribution with predicted differences in dormancy, lower left quadrant. **(D)** Frequency of ssDNA containing cell (EdU^+^ and/or BrdU^+^) in the indicated HSPC subsets. **(E)** Frequency of S-phase progression (BrdU^+^EdU^−^ as a fraction of EdU^+^ and/or BrdU^+^) in different HSPC populations (WT n=4, *Fancd2*^−/−^ n=5); \**P<0.05*.

A focused analysis of absolute FL HSC and HSPC (Lin^−^/Sca-1^+^/c-Kit^+^) (**Fig. 1B**) shows the predicted rapid gains in absolute number for WT and the lag for *Fancd2*^−/−^ KO cells during the critical HSC expansion phase from E12.5 – E14.5.

We reasoned that in spite of numerical HSC equivalency between genotypes at E12.5 and in the adult BM, experimental proliferative stress should reveal the same functional deficits in *Fancd2*^−/−^ seen at E13.5 and 14.5 in the fetal liver. Results in E12.5 and adult BM at 9 wk showed that significant clonogenic deficits emerge among colony forming cells, including the most primitive *Fancd2*^−/−^ subset (CFU-GEMM) (**Fig. S1C**). To assess *in vivo* repopulation capacity, we performed serial transplants of E12.5 *Fancd2*^−/−^ FL cells that confirmed deficits following the replicative stress of repopulation, which occurs with increasing p53 phosphorylation of HSC (**Fig. S1DE**). Intriguingly, with secondary transplantation, *Fancd2*^−/−^ cells showed low chimerism and revealed a peripheral blood myeloid (Mac1, Gr-1) bias, that phenocopies the behavior at steady state of myeloid progenitors (Fig. S1F).

These experiments suggest and confirm that FA HSC at E12.5 are vulnerable, but the physiologic functional deficits in FA HSC only emerge after E12.5, in response to proliferative cues in the FL.

### Fancd2 deletion causes S-phase entering delay in FL HSC

A range of cell cycle abnormalities has been described in FA, with prominent loss of quiescence and adult HSC exhaustion (Brosh Jr et al., 2017; Nalepa and Clapp, 2018) judged by Ki-67 staining for G0/G1. With roughly 90% of fetal HSC typically in cycle in the WT FL before transition to a more quiescent phenotype postnatally (Copley et al., 2013), we observed that fetal FA FL HSC are in fact hypoproliferative and fail to adequately expand, even as they positively stain for Ki67 (Domenech et al., 2018; Yoon et al., 2016). To resolve the conflicting observations, we undertook detailed studies using a timed 5-ethyl-2’-deoxiuridine/Bromodeoxyuridine (EdU/BrdU) sequential injection into pregnant E13.5 dams (Akinduro et al., 2018). Conceptually, cells entering S-phase during the two hours following EdU injection will initially become EdU^+^. This is followed by BrdU injection, when cells newly entering S-phase become EdU^−^/ BrdU^+^, cells exiting S-phase become EdU^+^/ BrdU^−^, and those that continue to remain in S-phase stain double positive EdU^+^/ BrdU^+^ (**Fig. 1C**). Our results show FL HSC enter S-phase more frequently than BM HSC, and a significantly greater fraction of *Fancd2*^−/−^ FL HSC and HSPC stain EdU^+^ and/or BrdU^+^ compared to WT (**Fig. 1D**), indicating an increased population of cells containing newly replicating single strand DNA (ssDNA). This is consistent with an increase in ssDNA observed by alkaline comet assay we previously reported (Yoon et al., 2016). Concurrently, the frequency of EdU^−^/ BrdU^+^ HSC and Lin^−^ cells was decreased in *Fancd2*^−/−^ compared to WT (**Fig. 2E**), indicative of delayed S-phase entry. These results confirm *Fancd2*^−/−^ FL HSC as less quiescent, but demonstrate a failure to progress through S-phase.

**Figure 2.**
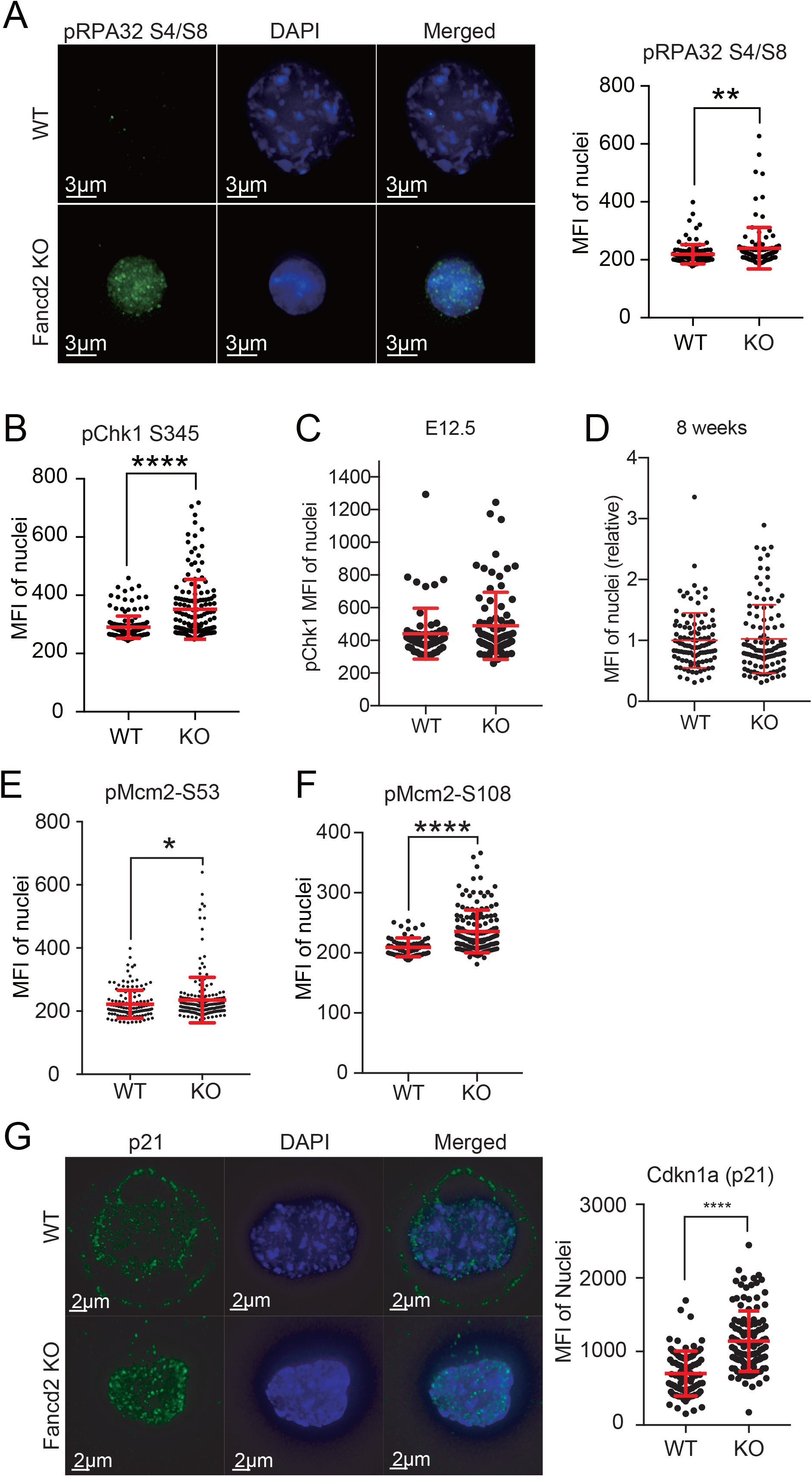
Replication stress response was detected in Fancd2^−/−^ FL HSPC. **(A-E)** Immunofluorescence (IF) of **(A)** pRPA32 S4/S8 (WT: n=5 pups, 116 cells, Fancd2^−/−^: n=4 pups, 111 cells), **(B)** pChk1 S345 (WT: n=9 pups, 175 cells, Fancd2^−/−^: n=4 pups, 146 cells), **(C)** pMCM S53 (WT: n=7 pups, 140 cells, Fancd2^−/−^: n=5 pups, 197 cells), **(D)** pMCM S108 (WT: n=4 pups, 64 cells, Fancd2^−/−^: n=6 pups, 149 cells), **(E)** Cdkn1a(p21) (WT n=4 pups, 84 cells, Fancd2^−/−^; n=4 pups, 120 cells). All data are measured mean fluorescent intensity (MFI) of nuclear in E13.5 FL WT and Fancd2^−/−^ HSPC. **(F, G)** IF of nuclear localized pChk1 S345 in **(F)** E12.5 HSC (WT n=2 pups, 63 cells, Fancd2^−/−^; n=5 pups, 81 cells) and **(G)** 8 weeks adult BM HSPC (used relative expression, WT n=2 pups, 106 cells, Fancd2^−/−^; n=2 pups, 113 cells).; *P<0.05, **P<0.01, ***P<0.001***P<0.0001.

### Replication stress is increased in the Fancd2 deficient FL HSC

FANCD2 specifically cooperates in sensing and resolving replication stress following experimental induction with hydroxyurea, aphidicoline and other agents. We hypothesized that the physiologic role of FANCD2 *in vivo* is to counter replication stress to rapid rates of proliferation in the fetal HSC pool. To determine whether *Fancd2*^−/−^ HSC experience replication stress, we checked replication associated protein (RPA)32; known to rapidly stabilize nuclear ssDNA during stalled replication. Our data in *Fancd2*^−/−^ HSPC showed increased phosphorylation of RPA32 with unchanged RPA70 protein levels (**Fig. 2A and S2A**). Phosphorylation of RPA-ATR is typically followed by phosphorylation of Chk1 (pChk1), which we found to be significantly increased as well (**Fig. 2B**). We reasoned that slower division cycles in adult phenotype (9 weeks) and pre-expansion E12.5 FL HSPC would fail to increase pChk1, and we found no significant differences between WT and *Fancd2*^−/−^ (**Fig. 2CD**). These results reflect the unique replication stress conditions during FL HSC expansion and the vulnerability of FA cells at this stage.

Deficits in mini chromosome maintenance (MCM) proteins are associated with replication stress and cell cycle defects in aging HSC (Flach et al., 2014). When we investigated activity in FA phenotype FL cells, we observed significantly increased phosphorylation of MCM2, a marker of replication fork stalling, at both S53 and S108 sites in *Fancd2*^−/−^ fetal HSPC (**Fig. 2EF**). Unlike the situation in aging HSC, we observed no evidence of broad transcriptional dysregulation when we conducted a gene expression survey of the additional DNA replication licensing factors MCM 2-7,Cdc1-7 kinases, and its activation co-factor Dbf4. Only Cdc7 was transcriptionally increased; however, there was no change at the protein level (**Fig S2BC**).

Transcriptional activity of CDKN1A(p21) in FA HSPC is typically considered a consequence of p53 pathway activation. On the other hand, p21 nuclear localization is a p53-independent event and associated with replication fork stalling (Karimian et al., 2016; Pietras et al., 2011; Li et al., 1996; Ma et al., 2013). Results from our studies of FL HSPC demonstrate a significant increase in nuclear CDKN1A(p21) in *Fancd2*^−/−^ compared with WT **(Fig. 2G**). Moreover, we considered the potential increase in RNA/DNA hybrids (termed R-loops), increased under experimentally induced replication stress. Unexpectedly, R-loop foci in *Fancd2^^−/−^^* KSL cells were decreased compared to WT (**Fig. S2D)**. Altogether, the data further show that *Fancd2*^−/−^ HSC experience exacerbated replication stress in the FL.

Cells completing replication under stress conditions can convert ssDNA breaks to dsDNA breaks, and other investigators have found low levels of γ-H2AX lesions without experimental induction. However, we examined fetal *Fancd2*^−/−^ HSPC and observed no spontaneous increase in γ-H2AX intensity, consistent with an absence of apoptosis markers and a lower rate of cell cycle completion (**Fig. S3F;** Yoon et al., 2016; Suzuki et al., 2016; Domenech et al., 2018). The aggregate data suggest that replication stress is a plausible cause for the observed delays in S-phase progression and deficits in FA FL HSPC expansion.

### Tgf-*β* inhibition rescues CDKN1A (p21) activation and Fancd2^−/−^ FL progenitor formation but not pChk1 activation

To address how CDKN1A(p21) activity is regulated in fetal *Fancd2*^−/−^ HSC, we considered Tgf-*β* signaling, a potent p53-independent activator of Cdkn1a(p21) in proliferating cells (Datto et al., 1995), and recently shown to be constitutively active in adult *Fancd2*^−/−^ HSC (Ceccaldi et al., 2012; Karimian et al., 2016). First, we measured Tgf-*β* receptor 1 (Tgfbr1) expression and found it to be increased in *Fancd2*^−/−^ HSC (**Fig. 3A**). Intriguingly, we show that pharmacological inhibition of Tgf-*β* by SD208 reverses the nuclear localization of Cdkn1a(p21) (**Fig. 3B**) and rescues the clonogenicity of primitive CFU-GEMM myeloid progenitors (**Fig. 3C**). However, in agreement with the notion of differential regulation for S-phase entry *versus* cell cycle progression (Rodriguez and Meuth, 2006), SD208, did not reverse pChk1 activation (**Fig. 3D**). These observations may suggest that the coincident increase observed in pChk1 after inhibition of Tgf-*β* represents as an aggravated replication stress response following release from Cdkn1a(p21) cell cycle inhibition.

**Figure 3.**
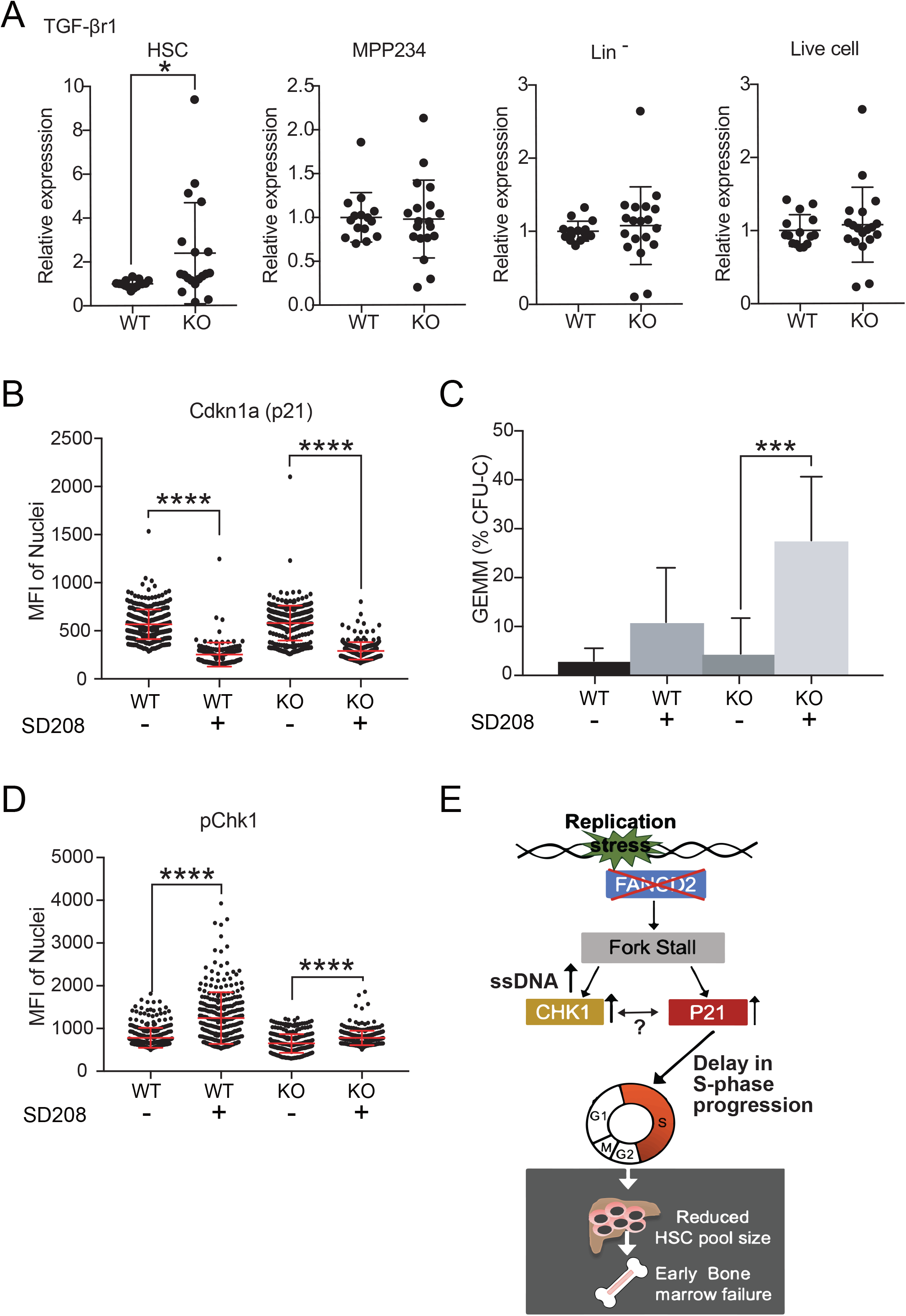
Tgf-*β* inhibition alters Cdkn1a(p21) localization, clonogenicity and pChk1 activity. (**A**) Evaluate Protein Tgfbr1 volume in HSC, MPP234, Lin^−^ and Total WT and Fancd2^−/−^ by FACS (WT; n=12, Fancd2^−/−^; n=19, E13.5, *\*P*<0.05). (**B**) IF of Cdkn1a(p21) after *in vitro* culture 48h with SD208 10uM treatment of E13.5 FL WT and Fancd2^−/−^ HSPC: (WT: n=6 pups, 338 cells, WT+SD208: n=6, 132 cells, Fancd2^−/−^: n=3 pups, 333 cells Fancd2^−/−^+SD208: n=3 pups, 212 cells; *\*\*\*\*P*<0.0001). (**C**) CFU assay and counted d14 GEMM colony frequency of E13.5 FL WT and Fancd2^−/−^ with and without SD208 (WT: n=3, WT+SD208 n=3, Fancd2^−/−^: n=9, Fancd2^−/−^+SD208 n=9; *\*\*\*P*<0.001). **(D)** IF of pChk1 S345 after 48h of in vitro culture of E13.5 FL HSPC WT and Fancd2^−/−^ with and without SD208 (WT: n=6 pups, 357 cells, WT+SD208: n=4, 276 cells, Fancd2^−/−^: n=3 pups, 320 cells Fancd2^−/−^+SD208: n=3 pups, 318 cells; *\*\*\*\*P*<0.0001).

## DISCUSSION

Experimental manipulation in murine models of FA provides strong evidence that multiple, non-mutually exclusive mechanisms contribute to the p53-dependent pro-apoptotic phenotype of adult FA HSPC that underlies the rapidly progressive decline in postnatal hematopoietic function in FA patients. We and others have previously reported the unexpected spontaneous deficits in the fetal HSPC pool of FA mice, and in this study, we identify a physiologic role for FANCD2 in an experimentally unprovoked *in vivo* system; namely, FANCD2 counters replication stress during rapid expansion in the fetal liver to attain proper HSC pool size and sustain lifelong hematopoietic needs (**Fig. 3E**). Such a role is consistent with HSC exhaustion that follows proliferative stress after experimental poly-I:C injection or serial transplantation to in FA deficient animals (Walter et al., 2015) and validates *in vitro* studies of the role FA proteins play in the replication stress response (Schlacher et al., 2011; Schlacher et al., 2012; Thompson et al., 2017).

The lack of fetal expansion was not explained by apoptosis, as seen in adult FA HSC. Moreover, such a hypoproliferative phenotype was difficult to reconcile with Ki67 cell cycle staining and we opted for greater cell cycle phase resolution using sequential EdU/ BrdU injections. These studies indeed confirmed that delays in S-phase entry and progression prevent fetal HSC pool expansion in *Fancd2*^−/−^. This ioffers a plausible explanation for the fetal FA HSC phenotype, given most HSPC are in cycle, than the mere loss of quiescence that explains exhaustion in adult FA deficient HSC (Copley et al., 2013).

Mechanistically, FANCD2 serves as a histone chaperone and may regulate accessibility of both replication- and transcription complex proteins to DNA (Sato et al., 2012). As a result, FANCD2 effectively suppresses the recruitment of additional replication origins during experimentally induced replication stress (Chaudhury et al., 2013; Thompson et al., 2017), consistent with our observed increase in replicated ssDNA and newly synthesized RNA (Fig. 1D, data not shown). Phosphorylation of MCM2 at both S53 and S108 sites in *Fancd*^−/−^ FL HSC further confirm that FA proteins also counter replication fork stalling *in vivo*.

Phosphorylation of RPA and Chk1 are canonical events during the replication stress response, and are significantly increased in fetal FA HSC during expansion in the liver (Fig. 2; Zeman and Cimprich, 2014). Importantly, pChk1 activation is not seen in E12.5 and adult BM HSPCs, indicating unique proliferative pressure in the FL during HSC expansion, and provides the physiological context for replication stress whereby Cdkn1a(p21) activity at the S-phase transition effectively restrains HSC proliferation during a critical developmental window. Such a p53-independent function of Cdkn1a(p21) in limiting proliferation in the FA HSC pool is also supported by studies of human fibroblasts under experimental replication stress (Lossaint et al., 2013). In adult FA HSC, the increased expression of Cdkn1a(p21) is widely considered a consequence of p53 engagement (Ceccaldi et al., 2012; Zhang et al., 2016). However, the concurrent loss of FANCD2 and p53 function in mice conferred only partial HSC rescue, whereas the compound loss of Cdkn1a(p21) and FANCD2 (but intact p53) further aggravates FA HSC losses (Ceccaldi et al., 2012; Garaycoechea et al., 2018; Zhang et al., 2013). Along with our data, these observations support the existence of a model whereby there is a p53-independent mechanism of developmental HSC failure in FA that invokes such a role for Cdkn1a(p21).

We show *in vivo* differences in pChk1 and Cdkn1a(p21) between WT and *Fancd2*^−/−^ HSPC, that are not present in *in vitro* assays (Fig. **3B,D)**, and attribute this to differences in metabolism and cell cycle progression, as has been noted in previous studies (Beerman et al, *2014*).

Tgf-β is known to rescue clonogenicity and HSC deficits in adult FA BM cells by altering DNA repair pathway usage (Zhang, H et al, 2016). Our experiments show for the first time that SD208 (Tgf-*β* receptor1 inhibitor) treatment can also restore fetal FA multipotent colony formation and nuclear Cdkn1a(p21) localization. However, this does not resolve the underlying replication stress and actually aggravates pChk1 expression in *Fancd2*^−/−^ HSPC (Fig. 3). This may be relevant for the critical role Chk1 plays in maintaining self-renewal, safeguarding HSC pool integrity, and minimizing mutational burden (Schuler et al., 2019). As an important clinical corollary, the attenuated phosphorylation of Chk1, i.e. low CHK1 expression, seen in some adult FA patients leads to temporary improvement in hematopoietic function, that subsequently gives way to myelodysplastic clonal evolution (Ceccaldi et al., 2011). Thus, available evidence indicates that both Cdkn1a(p21) and pChk1 prevent apoptosis in fetal *Fancd2*^−/−^ HSPC. How they spatially and temporally interact in regulating cell cycle progression remains to be clarified.

In aggregate, our study reveals the origin and a physiologic mechanism for fetal hematopoietic failure in FANCD2 deficient animals, which may hold new therapeutic opportunity for the rescue of hematopoietic function in FA patients and offer insight for other BMF disorders.

## MATERIALS AND METHODS

### Animal husbandry and cells

C57/BL6 background Fancd2 KO mice (Houghtaling et al., 2003) were bred and used for experiments. WT and Fancd2 KO fetuses were harvested from timed pregnancies generated from crossing heterozygous female mice with heterozygous male mice WT and Fancd2 KO fetuses were harvested from timed pregnancies generated from crossing heterozygous female mice with heterozygous male mice. Fetal livers were harvested from pups and separated by mechanical disruption, filtration and subsequent red blood lysis to get mononuclear cells. Bone marrow was harvested from femur.

Animal husbandry, tissue harvest and processing were all described previously (Yoon et al., 2016). All animal experiments were approved by the OHSU and CHOP Animal Care and Use Committee.

### Immunophenotyping and FACS analysis

FL or BM mononuclear cells were stained with Lineage marker antibodies (CD3e, CD4, CD5, B220, Gr-1, Ter119) as well as HSPC markers: c-Kit, Sca-1, CD48 and CD150 for 30mins at 4 degree. Blocking and washing Buffer contained 2%FBS/ PBS. Dead cell exclusion staining used DAPI at 1ug/ml. Flow cytometric analysis was performed using FACS Canto2 and LSR2 instruments (Becton-Dickinson) as described (Yoon et al., 2016). Reagent Supplemental Table 1 for details.

### Colony Formation Unit (CFU) assay

Harvested and RBC lysed FL and BM were counted using Trypan blue stains and mixed with cytokine supplemented commercial mouse methylcellulose media (R&D Systems, HSC007), with aliquots divided into 3 × 3.5cm dishes, cultured at 37 degree. After 10-14 days, colony number and colony subtype were scored under an inverted light microscope.

### Serial Transplantation

Harvested and RBC-lysed E12.5 FL (5×10^5^) were injected via the tail vein in CD45.1 recipients that received 750cGy using a - irradiator (single dose). At 20 weeks from transplantation, animals were sacrificed, tissues analyzed and secondary transplantation was performed with injection of 1×10^6^ BM cells into 750Gy irradiated CD45.1 recipients. Peripheral blood from both primary and secondary recipients was analyzed for chimerism at 9 weeks from transplantation, using antibodies against Gr-1, Mac-1, B220, CD3e and DAPI, by FACS using a Canto2 (BectonDickinson).

### EdU/ BrdU cell cycle assay

We modified a previously reported assay **(Akinduro et al., 2018)** with sequential injection via the tail vein of E13.5 pregnant females (vaginal plug method) with 1mg of EdU, followed 2 hours later by 2mg of BrdU. After 30 minutes FLs were harvested and individually processed. Isolated FL mononuclear cells were stained with surface markers (CD150, CD48, c-Kit, Sca-1, Lin) and fixed in 2% paraformaldehyde (PFA) for 15 minutes, followed by permeabilization with 0.5% saponin and stained with anti-EdU-AF488. To stain with anti-BrdU antibody, we treated with 20ug of DNAse in PBS (containing Ca^++^, Mg^++^) at 37 degree for 40 minutes before staining with BrdU-AF647 (B35140, Thermo). Analysis was performed with FACS LSR2 (Becton-Dickinson).

### Analytical flow cytometry

FL or BM mononuclear cells were stained with surface markers and fixed with 2% PFA for 15 minutes, permeabilized with 0.5% saponin and stained anti-p53, anti-p53S15 and anti-Cdc7. Analysis was performed with FACS LSR2 (Becton-Dickinson) and data processed with Flowjo 10.5.0 to quantify mean fluorescent intensity (MFI). Supplemental Table 1 for Reagent details.

### Flow cytometric sorting

FL mononuclear cells were stained with CD150, CD48, c-Kit, Sca-1, Lin and DAPI (Thermo 62248, 1ug/ml) and sorted using an Influx Aria Fusion instrument (Becton-Dickinson).

### Immunofluorescence

Sorted cells (5-500 ×10^3^) were placed on glass slides using a cytocentrifuge, followed by incubation with or without Cytoskeletal (CSK) buffer for 10min, at room temperature and fixed in 4% PFA. Permeabilization was performed by 0.5% Triton and blocking was with 3% BSA/PBS at 37 degree for 30 minutes. Primary antibody staining was performed on parafilm at 37 degree for 30 mins, and secondary antibodies were used with 1:1000 dilution at 37 degree for 30min. For nuclear staining, DAPI was used at room temperature for 10min. For coverslip mounting we used Fluoromount-G (0100, Southern Biotech). Images were captured on a Core DV microscope (Olympus) and via LSR700 confocal microscopy (Carl Zeiss). Images were processed and analyzed with Imaris software (Bitplane). Supplemental Table 1 for Reagent details.

### Quantitative RT-PCR analysis

RNA from flow cytometrically sorted cells was isolated using the RNeasy mini and micro kit (QIAGEN). Reverse transcription was performed using SuperScript™ Master Mix (Invitrogen). For Q-PCR, we used FastStart Essential DNA Green Master (Roche) and LightCycer 96^®^ (Roche).

### Ex vivo cell culture

SLAM marker -sorted cells were placed in StemSpan (09650, Stem Cell Tech.) supplemented with 0.5% Penicillin/streptomycin, stem cell factor (250-03, Peprotech) 50ng/ml and thrombopoietin (315-14, Peprotech) at 50ng/ml.

### Quantification and Statistical Analysis

All numerical results were expressed as mean ± SD. Two-tailed Student’s t tests, Welch test and One-way ANOVA were performed for statistical analyses. All analyses were performed with GraphPad PRISM 7.0

## ACKNOWLEDGEMENTS

We are grateful for support by the Department of Pediatrics at OHSU and wish to thank Dr. Devo Goldman, Dr. Sherif Abdelhamed, John T. Butler and Dr. Qingshuo Zhang for support and guidance with select experiments. We acknowledge Yanet Wossenseged for her contributions to Figure 3E.

## Figure legends

**Figure S1.**
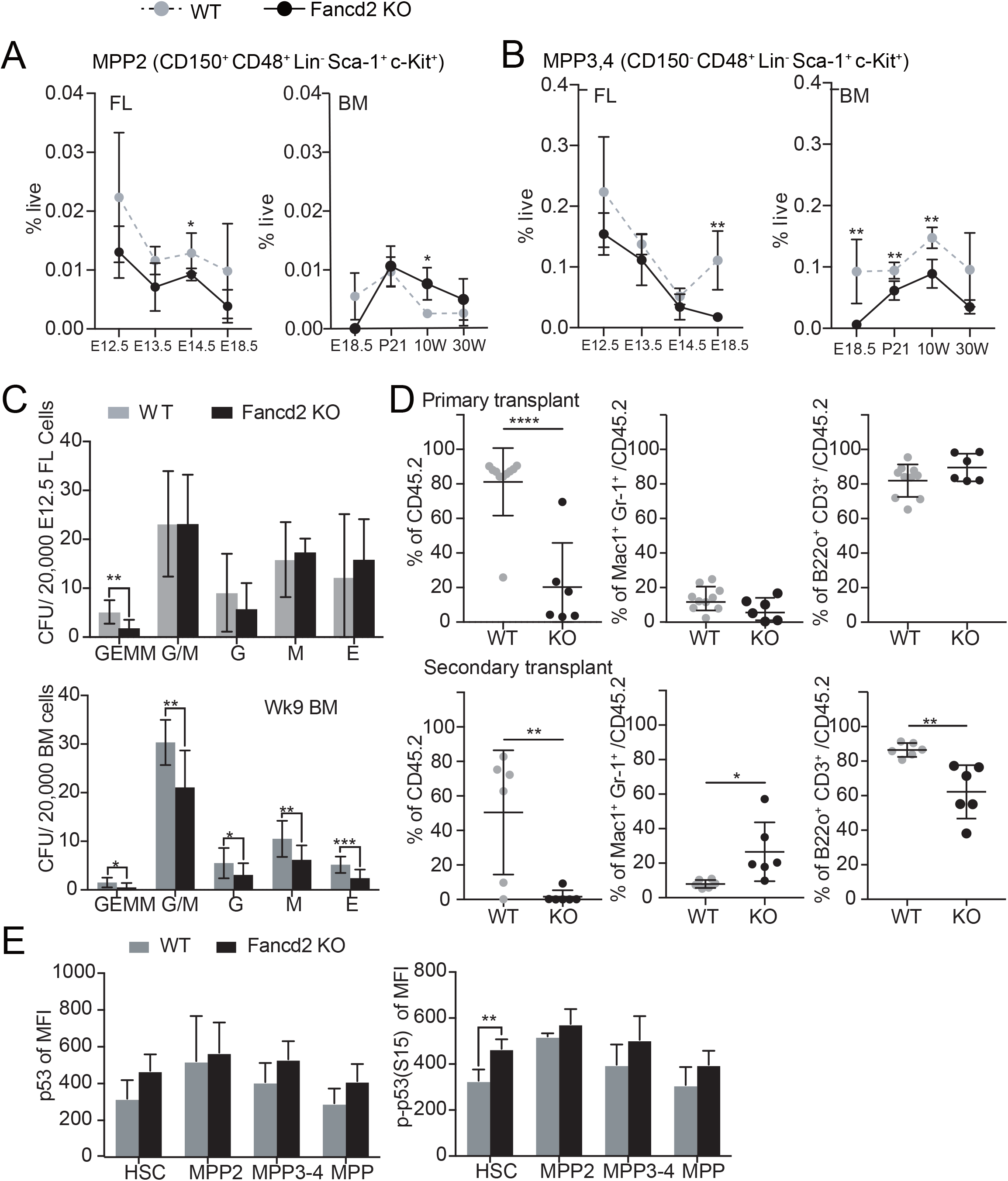
Fancd2 deficiency limits HSC expansion in the fetal liver. **(A,B)** Immunophenotyping was performed to determine the frequency of **(A)** CD150^+^ CD48^+^ LSK, MPP2 and **(B)** CD150^−^ CD48^+^ LSK, MPP3,4 **i**n WT and Fancd2^−/−^ cell across select time points in ontogeny. **A,B:** E12.5 n=3^+/+^, 7^−/−^, E13.5 n=4^+/+^, 5^−/−^, E14.5 n=9^+/+^, 6^−/−^, E18.5 n=7^+/+^, 3^−/−^, P21 n=5^+/+^, 6^−/−^, 10 Weeks(10W) n=4^+/+^, 4^−/−^, 30W n=3^+/+^,3^−/−^). **(C)** Colony formation from 20,000 E12.5 FL cells (upperpanel) and BM cells at 9 weeks of age (lower panel). **(D)** *In vivo* serial transplantation using 5×10^5^ E12.5 FL WT and Fancd2^−/−^ cells, showed peripheral blood chimerism of total(left panel), myeloid(mid panel) and lymphoid(right panel) in primary (9 weeks from transplantation, upper panels) and secondary transplantation (9 weeks from 2^nd^ transplantation, lower panels). *\*P*<0.05, *\*\*P*<0.01, *\*\*\*P*<0.001, *\*\*\*\*P*<0.0001 **(E)** p53(left panel) and p-p53 S15(right panel) protein volume at 9 weeks of HSC, MPP2, MPP3,4 and MPP total from transplantation of E12.5 FL WT and Fancd2^−/−^ (Fig. 1F) were detected by flowcytometry (WT; n=4, Fancd2^−/−^; n=4).

**Figure S2.**
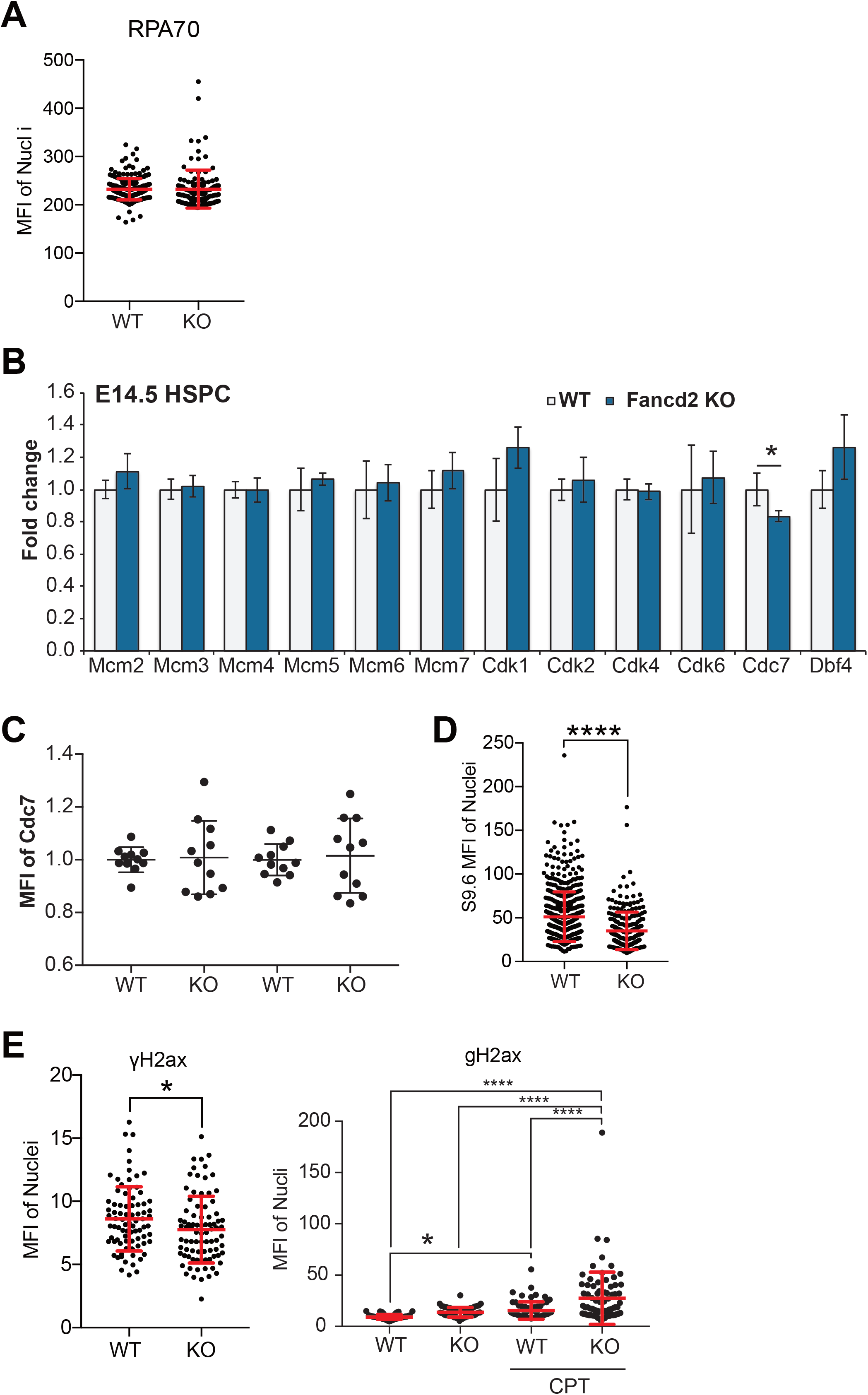
Fancd2^−/−^ FL HSC showed replication stress response. **(A)** IF of Rpa70 and measured nuclear localized MFI at E13.5 FL WT and Fancd2^−/−^ HSPC (WT; n=10 pups, 227 cells, Fancd2^−/−^; n=3 pups, 120 cells). **(B**) mRNA expression of Mcm2-7, Cdk1,2,4,6, Cdc7, Dbf4 of WT and Fancd2^−/−^ in E14.5 FL HSPC (E14.5 WT; n=4, Fancd2^−/−^; n=3). **(C)** Cdc7 protein volume expression in E13.5 WT and Fancd2^−/−^ HSC (WT; n=11, Fancd2^−/−^; n=11). **(D)** IF of S9.6 and measured nuclear localized MFI at E14.5 FL WT and Fancd2^−/−^ HSPC (WT; n=10 pups, 227 cells, Fancd2^−/−^; n=3 pups, 120 cells). **(E)** IF of γH2AX and measured nuclear localized MFI at E14.5 FL WT and Fancd2^−/−^ HSPC (left pannel; WT; n=2 pups, 83 cells, Fancd2^−/−^; n=2 pups, 84 cells) and ex vivo 1 hour cultured with or without CPT (50ug/ml) (right panel; WT; n=1 pup, 12 cells, Fancd2^−/−^; n=2 pups, 46 cells, WT+CPT; n=1 pup, 12 cells, Fancd2^−/−^+CPT; n=2 pups, 11 cells).

---

**Supplemental Table 1.**
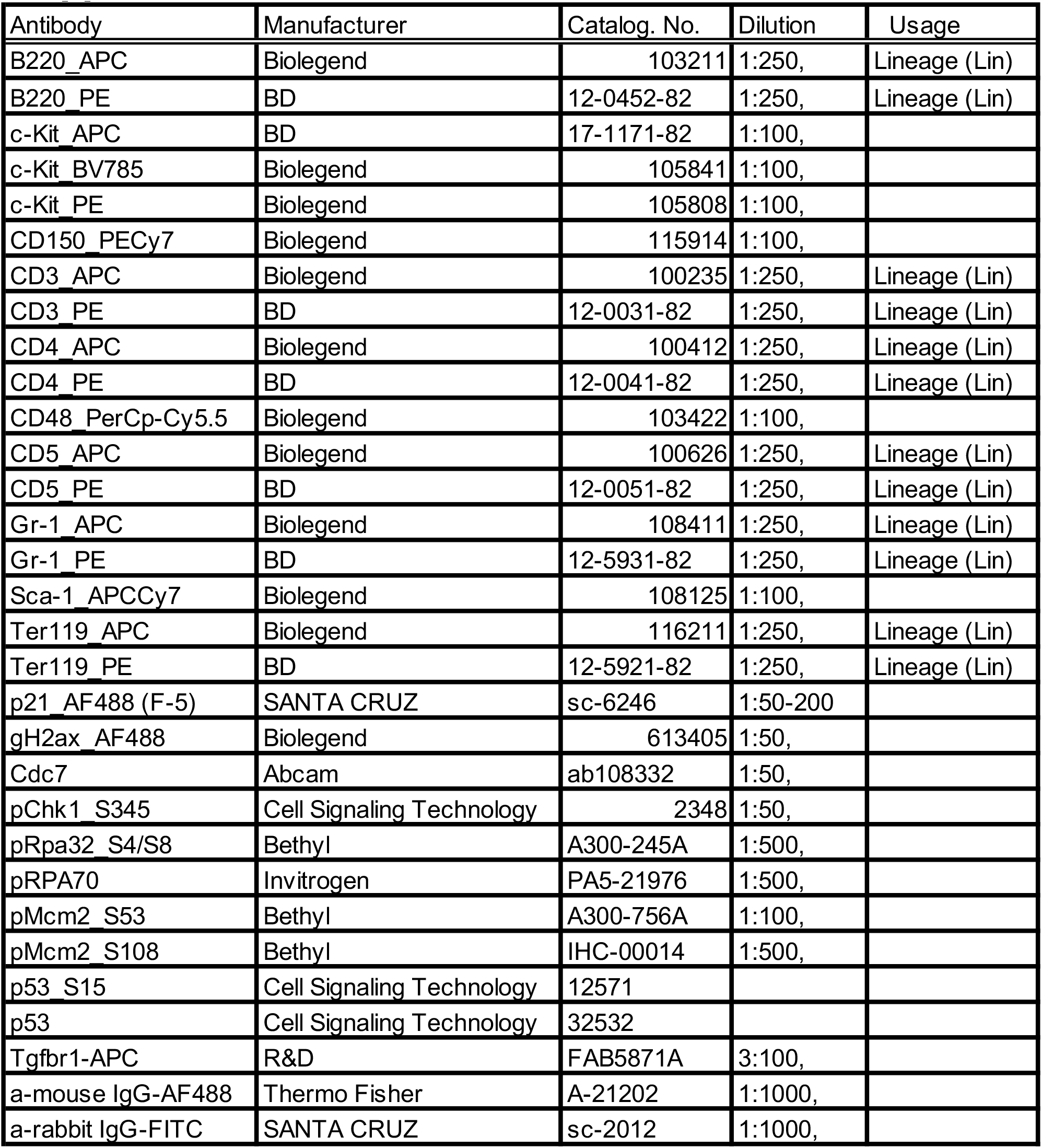

